# A pathogen-responsive gene cluster for the production of highly modified fatty acids in tomato

**DOI:** 10.1101/408518

**Authors:** Ju Eun Jeon, Jung-Gun Kim, Curt R. Fischer, Cosima Dufour-Schroif, Kimberly Wemmer, Mary Beth Mudgett, Elizabeth Sattely

## Abstract

In response to biotic stress, plants reshape their complement of lipids to produce suites of highly modified fatty acids that bear unusual chemical functionality. Despite their chemical complexity, proposed roles in pathogen defense and presence in crop plants, little is known about the biosynthesis of these decorated fatty acids. Falcarindiol is a prototypical member of a suite of acetylenic lipids from carrot, tomato, and celery that inhibits growth of several fungal strains and human cancer cell lines. Here we report a set of clustered genes in tomato (*Solanum lycopersicum*) that are required for the production of falcarindiol in leaves in response to treatment with an adapted fungal pathogen, *Cladosporium fulvum*. Our approach is based on correlation of untargeted transcriptomic and metabolomic data sets in order to rapidly identify a candidate biosynthetic pathway. By reconstituting the initial biosynthetic steps in a heterologous host (*Nicotiana benthamiana*) and generating stable transgenic pathway mutants in tomato, we demonstrate a direct role for three genes in the cluster in falcarindiol biosynthesis. This work reveals a mechanism by which plants sculpt their lipid pool in response to pathogens, and provides critical insight into the biochemistry of alkynyl lipid production.

**One Sentence Summary:** A biosynthetic gene cluster for the production of falcarindiol, a highly modified antifungal oxylipin found in edible plants.

## Main text

Modified lipids play a central role in plant defense (*1*, *2*). While structural lipids drawn from primary metabolism limit pathogen entry (e.g. extracellular cuticle), plants also reshape their complement of lipids in response to biotic stress to produce metabolites that function as signals or antimicrobial agents (*3*). A notable example is jasmonic acid, a ubiquitious plant hormone and oxylipin biosynthesized inducibly from membrane-derived linolenic acid that undergoes enzymatic tailoring (*4*, *5*). Acetylenic fatty acids are a related family of oxylipins present in seed oils or biosynthesized in response to pest and pathogen stress (*6*, *7*). These compounds are highly modified lipids widely distributed throughout the plant kingdom - many of which include unique sets of desaturations, acetylenic functionality, and oxidations (Fig. 1) (*8*).

**Fig 1.**
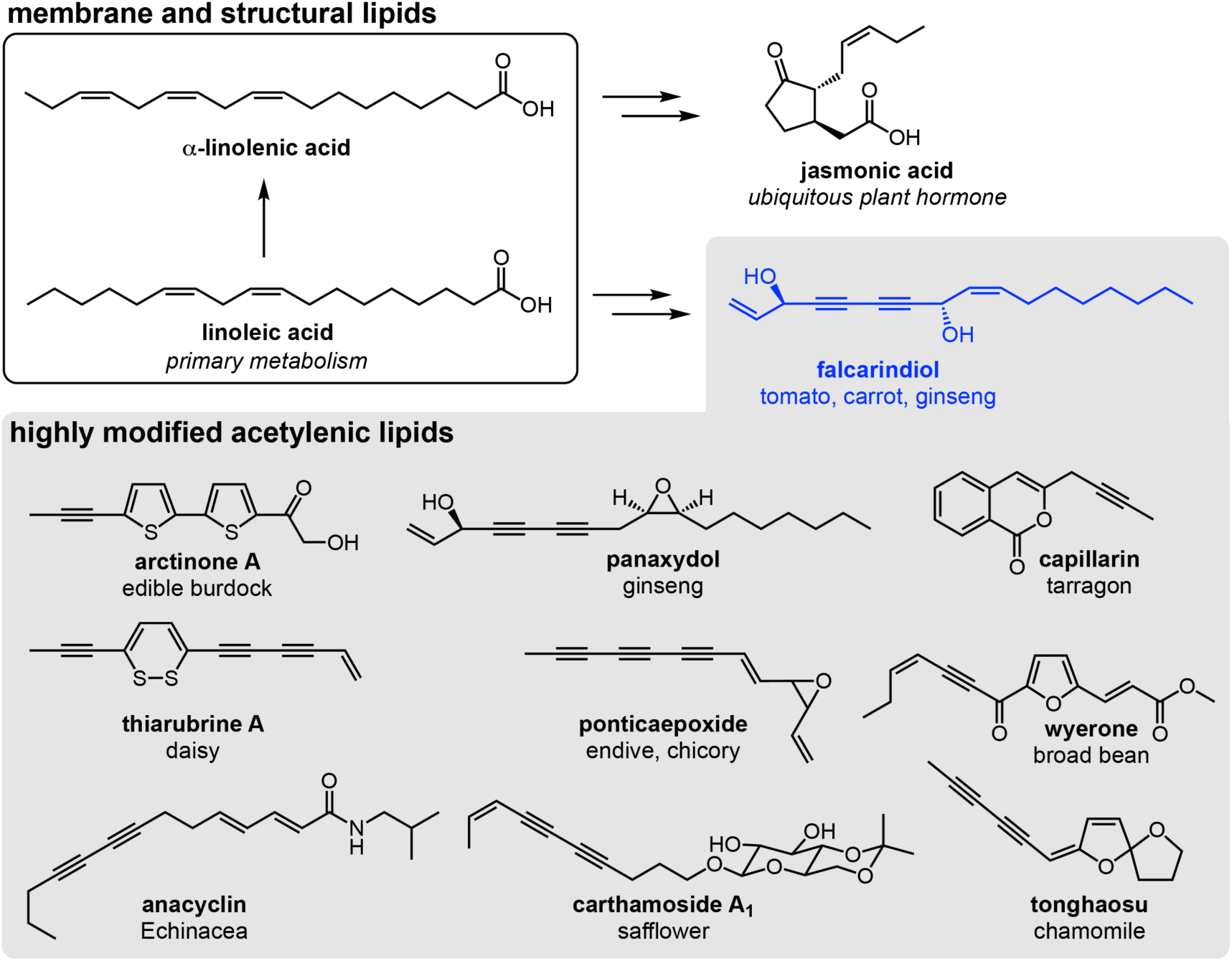
Representative acetylenic natural products found in edible plants. Linoleic acid produced in primary metabolism is released from plant membranes and is used in secondary metabolism to produce the hormone jasmonic acid. Linoleic acid is also a possible precursor for the synthesis highly modified acetylenic lipids. Falcarindiol is a representative acetylenic lipid found in tomato, carrot and ginseng, for which no biosynthetic genes have been described.

Compared to other broad families of plant metabolites (e.g. alkaloids and terpenes) (*9*), relatively little is known about the biosynthesis of highly modified fatty acids (*10*), limiting our ability to take advantage of the biosynthetic capacity of plants to make valuable lipids through metabolic engineering (*11*). As part of an effort to unravel the biosynthetic logic that governs the production of modified lipids, we chose to investigate the biosynthesis of falcarindiol, a representative oxylipin and odd-chain alkane found in edible plants that bears conjugated acetylenic functionality, oxidation, and an unusual terminal vinyl group (*12*, *13*). In addition to its chemical novelty, falcarindiol is a dietary metabolite with potent antifungal activity and cytotoxicity against several cancer cell lines (*14*). Despite the demonstrated biological activity of falcarindiol and its presence in edible crops - abundant in carrot (6-60 mg/kg fresh weight in root tissue) (*15*) and ginseng root and induced by fungal pathogens in the leaves and fruit of tomato (*16*) - no genes have been associated with the biosynthetic pathway. We predicted that the chemical modifications to the fatty acid backbone of falcarindiol likely involve a series of atypical transformations catalyzed by classical Fe-S desaturases and/or heme-containing oxidases that have evolved novel function (*17*). Intriguingly, one such enzyme has been described in *Crepis alpina*: Crep1, a novel acetylenase that installs an alkyne in linoleic acid to generate crepenynic acid that accumulates in the seed oil of this plant (*18*). However, no homolog in other plants has been characterized or associated with the biosynthesis of more highly modified lipids.

In order to uncover candidate genes in tomato for the biosynthesis of falcarindiol, we leveraged the observation that falcarindiol accumulation can be induced in tomato plants by biotic elicitors. Reasoning that that temporal changes in metabolite levels would likely correlate with transcript levels for biosynthetic enzymes, we used a combination of untargeted metabolomics and RNA sequencing on paired samples of tomato leaf tissue challenged with biotic stress. To capture a wide range of pathway expression levels, we challenged tomato leaves with a diverse panel of microbial elicitors over a time course of 48 hours (See fig. S1 and Table S1 for experimental design). We chose a set of elicitors that would likely invoke varied effects on pathway expression, including two microbe-associated molecular patterns (MAMPs) (*19*) (fungal chitin and bacterial flagellin, i.e. Flg22), two plant-associated pathogens (*Cladosporium fulvum* and *Xanthomonas euvesicatoria*), and three human-associated microbes (*Malassezia restricta*, *Staphylococcus epidermidis* and *Propionibacterium acnes*), for a total of 7 treatments and two mock controls (for fungal and bacterial elicitors). We reasoned that adapted plant-associated microbes (microbes that have evolved with their natural hosts) and non-adapted microbes may elicit distinct and shared responses by the plant and would allow us to capture changes in the expression of a broad set of genes that contribute generally to defense and specifically to falcarindiol biosynthesis.

Quantitative metabolomic profiling using LC-MS analysis revealed significant and selective changes in metabolite levels including falcarindiol in response to elicitor type (fig. S2 and S3A), suggesting the utility of our data set for pathway discovery. For example, falcarindiol production is reproducibly induced only by a subset of elicitation conditions (Fig. 3A); *C. fulvum*, *M. restricta*, and *S. epidermidis* elicited tomato leaves after 12, 24, and 48 hours post-infiltration). By contrast, mass signatures corresponding to hydroxy cinnamic amides (*20*) and compounds related to the sterol alkaloid tomatidine (*21*) varied in abundance across the sample panel (fig. S3B-G), and are notably different from the patterns of falcarindiol accumulation. This suggested to us that metabolite-transcript correlation analysis for falcarindiol could help pinpoint candidate pathway genes that install the acetylenic functionality and provide a starting point for elucidating the genetic basis for the biosynthetic pathway.

Based on previous examples of plant fatty acid metabolic pathways and characterized acetylenases (*1*, *22*), we formulated a hypothesis for falcarindiol biosynthesis that involves many steps carried out by desaturases (for proposed pathway, see fig. S4). We proposed that an early step in the pathway might involve conversion of linoleic acid to crepenynic acid via a Crep1-like acetylenase (*18*). Crep1 is one of the few enzymes known to be involved in modified fatty acid biosynthesis; therefore, we hypothesized that a related acetylenase in tomato is likely required for shunting fatty acids from primary metabolism into secondary metabolic pathways. Given that only a few amino acid changes distinguish acetylenases from other desaturases (*17*) we could not identify a clear ortholog of Crep1 in tomato. Instead, we used a more general search for desaturases to generate a list of candidate acetylenases.

Of 59 unique desaturases predicted in the Heinz tomato genome (*23*) (Heinz-SL2.50 version, Phytozome database) using a combination of BLAST and Pfam annotation based on the *Arabidopsis thaliana* proteome (Table S3), we detected 40 of the corresponding transcripts in our RNA-Seq dataset. We correlated the expression level of these transcripts with our LC-MS-based measurements of falcarindiol accumulation (Fig. 2A & 2B) to prioritize the putative desaturases likely involved in the installation of the falcarindiol alkynes. Unexpectedly, we noted that of the six genes whose expression levels correlate best with falcarindiol accumulation (based on *r*-value from linear regression analysis, cut-off value > 0.2) (fig. S5), four were located sequentially on chromosome 12 in the same 20 kb region of the genome (Fig. 2C, Solyc12g100240-Solyc12g100270) and highly co-expressed (fig. S6). Three of the genes are annotated as desaturases/acetylenases and the fourth as a decarbonylase, suggesting that this locus encodes a cluster of enzymes associated with fatty acid metabolism. The presence of a decarbonylase was notable given that falcarindiol lacks the carboxylic acid typical of a fatty acid. Biosynthetic gene clusters have been reported for the production of many classes of plant metabolites (*24*), but only one has previously been reported for the production of a polyketide (*25*).

**Fig 2.**
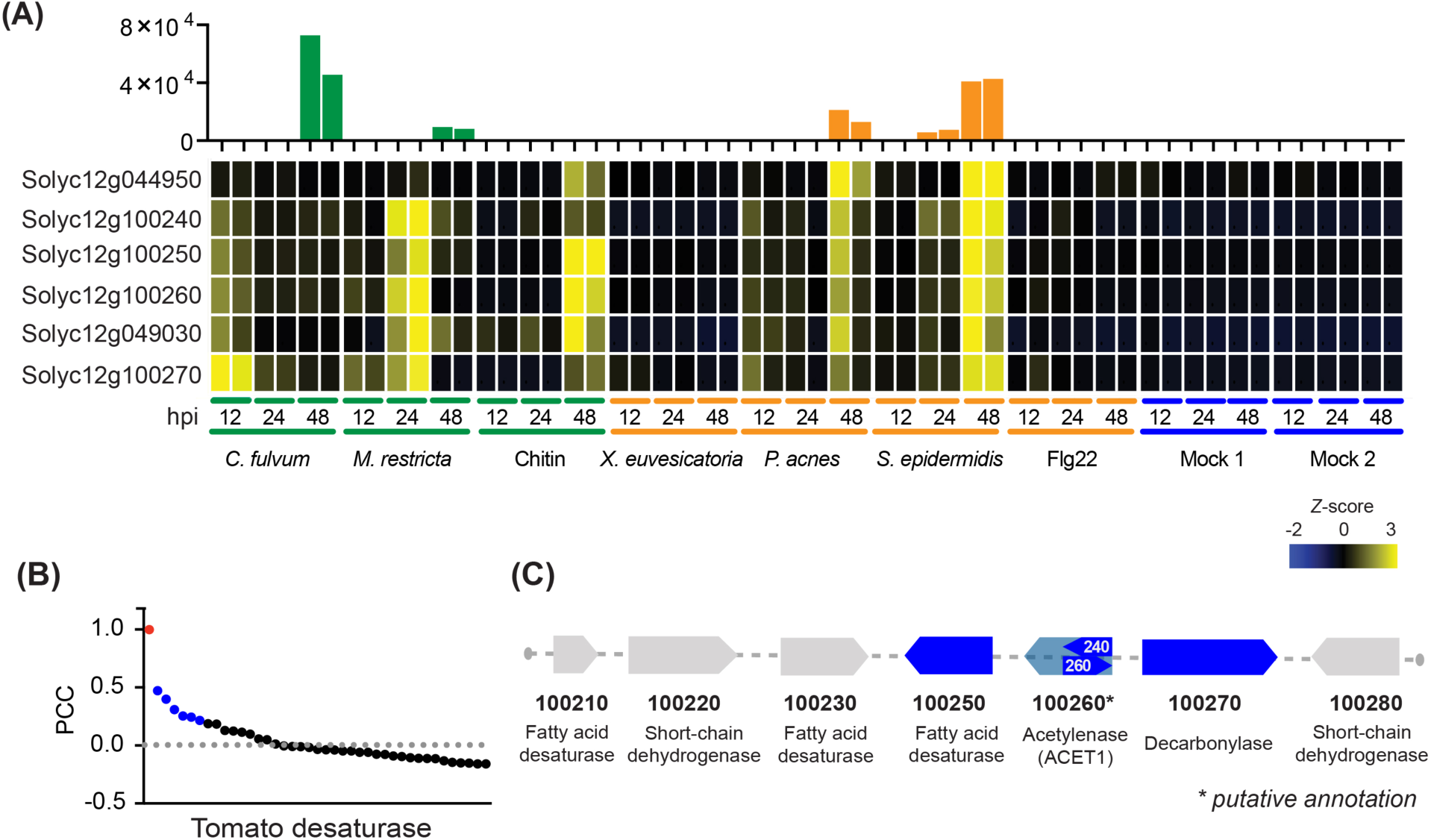
Identification of putative non-heme diiron enzymes including desaturases and decarbonylases to discover biosynthetic genes involved in falcarindiol production. (A) Correlation of falcarindiol levels (bar graph shows extracted ion chromatogram (EIC) peak integration (m/z 283.16, [M+Na]^+^) using LC-MS) to transcript abundance of six candidate acetylenase genes (heatmap illustrates Z-score determined using counts per million base pairs values obtained using RNA-Seq) in tomato leaves under various elicitation conditions. Transcripts are sorted by their Pearson’s *correlation* coefficient (PCC) values against falcarindiol production (SI SI5). Colors represent elicitor types (green: fungal elicitor, orange: bacterial elicitor, blue: Mock, water). Experiments were conducted in two separate batches: Mock 1 for MAMPs and human-associated microbes; Mock 2 for plant-associated microbes. (B) PCC values for 44 candidate tomato desaturases’ transcript abundance (CPM) against falcarindiol levels (red: falcarindiol, blue: six candidate genes in Fig. 2(A), black: remaining 38 desaturases detected in RNA-Seq analysis). (C) Genomic organization of the candidate metabolic gene cluster (genes related to the falcarindiol pathway are highlighted in blue and 100260 represents *de novo* assembled transcript). Genes are denoted without ‘Solyc’ and chromosome number.

Two of the candidate genes in the cluster, Solyc12g100240 and Solyc12g100260, have overlapping identical sequence (Fig. 2C) and appear to encode a partial desaturase. We suspected that these fragments represent an incomplete gene sequence due to misannotation of the Heinz tomato genome. We performed *de novo* assembly of our RNA-Seq data from VF36 tomato to examine gene structure at this locus. This analysis revealed a single transcript (1140 nts) that contains both the Solyc12g100240 and Solyc12g100260 sequences and is predicted to encode a 380 kDa protein characteristic of a typical desaturase. Notably, an identical transcript was detected in a Heinz cDNA library, corroborating the new gene sequence assignment. We now refer to this gene as Solyc12g100260 (Fig. 2C) (*26*).

In order to determine if any of the genes in the cluster possess acetylenase activity, we overexpressed the enzymes in *Nicotiana benthamiana* using *Agrobacterium*-mediated transient expression. Three days post infiltration, *N. benthamiana* leaves were collected and saponified methanol extracts were analyzed by LC-MS for accumulation of crepenynic acid. The *m/z* value for the crepenynic acid proton adduct was only detected in *N. benthamiana* tissue expressing the cDNA for Solyc12g100260 (Fig. 3). The adduct exhibited the same retention time as the control product from *Crep1* expression (fig. S7) and the presence of an alkyne was supported chemical derivatization of the alkyne using ruthenium-catalyzed azide-alkyne cycloaddition click chemistry (fig. S8-10) (*27*). These data indicate that Solyc12g100260 (renamed *ACETYLENASE1* or *ACET1*) encodes an acetylenase that converts linoleic acid to crepenynic acid, suggesting it catalyzes the first step in the falcarindiol pathway in tomato. Remarkably, reverse correlation analysis using *ACET1* gene expression as bait against the collection of all metabolites detected in tomato leaf extracts after elicitation revealed falcarindiol as the metabolite whose production is most correlated with *ACET1* expression (fig. S11). This analysis highlights the power of untargeted correlation of gene expression to metabolite production for pathway discovery, and corroborates the proposed role of *ACET1* in falcarindiol biosynthesis.

**Fig 3.**
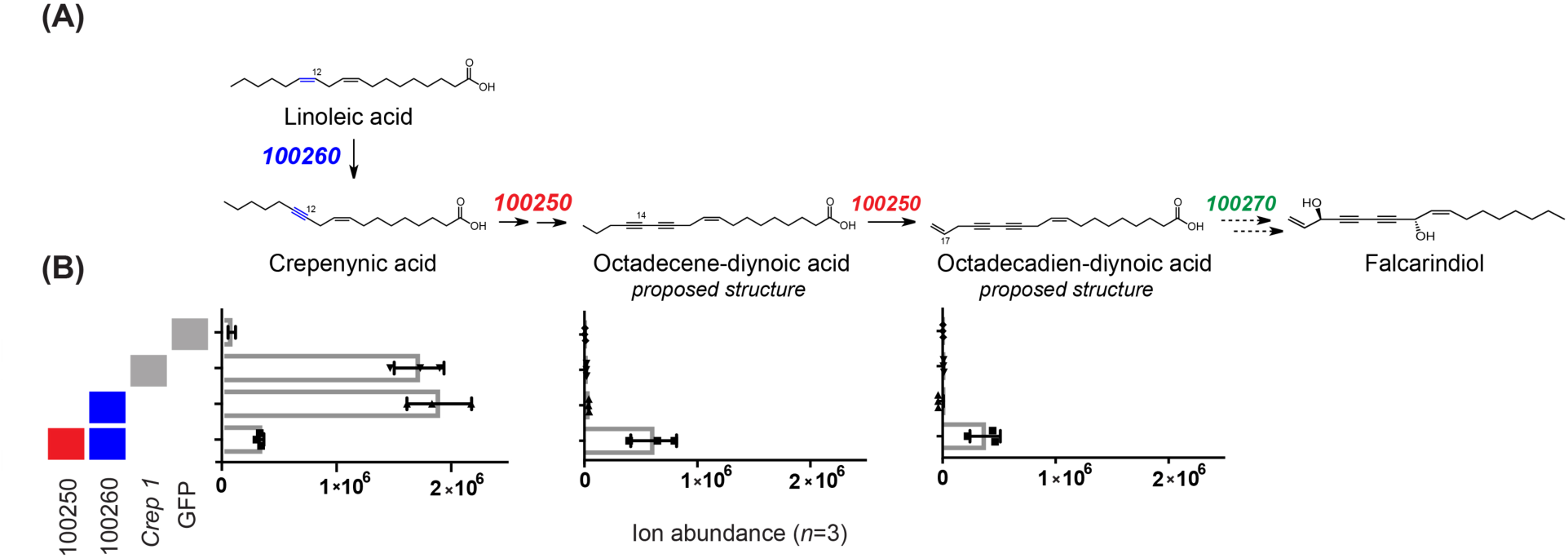
Reconstitution of tomato biosynthetic genes for highly modified fatty acid in *Nicotiana benthamiana*. (A) Proposed pathway from linoleic acid to falcarindiol and role for candidate genes from identified gene cluster. Genes are denoted without ‘Solyc’ and chromosome number. Proposed chemical structures of octadecene-diynoic acid and octadecadien-diynoic acid based on LC-MS data and acetylene derivatization using click chemistry. (B) Metabolite accumulation *in N. benthamiana* leaves determined by integration of EIC for indicated compound (ion abundance plotted as mean ± SD (*n = 3*). Grey and colored boxes represent transiently expressed enzymes indicated below (100250, 100260, Crep1 from *Crepis alpina* or green fluorescent protein (GFP) as negative control).

Further inspection of the gene cluster suggested that in addition to *ACET1,* Solyc12g100250 and Solyc1210070 might be involved in falcarindiol biosynthesis. Not only are their annotated enzymatic activities consistent with the proposed biosynthetic pathway (e.g. further desaturation and decarbonylation), but Solyc12g100250 and Solyc12g100270 are the genes whose expression levels correlate best with those of *ACET1* (Fig. S5). To test a role for these genes in falcarindiol biosynthesis, we transiently co-expressed each with *ACET1* using *Agrobacterium*- mediated expression in *N. benthamiana* to identify potential modified fatty acid metabolites representing downstream intermediates in the pathway. Untargeted metabolite analysis in leaves expressing both Solyc12g100250 and *ACET1* revealed depletion of crepenynic acid and accumulation of two new mass signatures whose formulae correspond to octadecene-diynoic acid and a compound with an additional desaturation, octadecadiene-diynoic acid (Fig. 3, fig. S9). Although we have not been able to confirm the proposed structures illustrated in Fig. 3 using authentic standards, MS/MS analysis and derivatization of the alkyne using the aforementioned click chemistry (fig. S9-10) suggest they are modified fatty acids that contain acetylenic functionality. These data provide biochemical evidence that Solyc12g100250 encodes a desaturase that can use crepenynic acid as a substrate. It is unclear if Solyc12g100250 is responsible for multiple steps or functions with ACET1 to produce the detected molecules.

Solyc12g100270 is one of five tomato homologs of CER1, a fatty acid decarbonylase in *A. thaliana* which functions with its partner enzyme CER3 to catalyze very-long chain fatty acid decarboxylation to produce an alkane in wax biosynthesis (*28*). Notably, the CER1/CER3 pair provides precedent for a biosynthetic route to modified plant lipids with odd chain lengths. Given that Solyc12g100270 is homologous to CER1, we hypothesized that this enzyme might convert the carboxylic acid of a falcarindiol precursor to the alkane terminus, which is a signature for this modified fatty acid (fig. S4). However, transient expression of Solyc12g100270 with or without Solyc12g100250, ACET1, and *Arabidopsis* CER3 in *N. benthamiana* did not lead to the production of a new metabolite. It is possible that Solyc12g100270 is not expressed in an active form in *N. benthamiana*, is missing a partner protein, or acts further downstream in the pathway.

As an alternative approach to study the role of Solyc12g100270, we tested directly if the gene is required for falcarindiol production using CRISPR/Cas9 genome editing in tomato (*29*). In addition, we targeted *ACET1* (Solyc12g100260) in order to validate the first step in the pathway. Multiple knock-out mutants (designated *ΔACET1* and Δ270) were generated in the tomato VF36 background (Fig. 4A, fig. S12-S13). Untargeted metabolite analysis of tomato leaf extracts from WT and multiple independent *ΔACET1* and Δ270 mutant lines treated with *C. fulvum* revealed that falcarindiol is among three major metabolites present in WT but absent in both biosynthetic mutants (Fig. 4B, Fig. S14). Proposed formulae for the other two mass signatures are consistent with modified fatty acid scaffolds, although we have not yet been able to confirm their chemical structures (Fig. S14). These data corroborate the biochemical activity observed for *ACET1* in *N. benthamiana* and indicate that both *ACET1* and Solyc12g100270 are required for production of falcarindiol in tomato. Furthermore, our results provide direct support for the role of the *ACET1* gene cluster for the synthesis of modified fatty acids.

**Fig 4.**
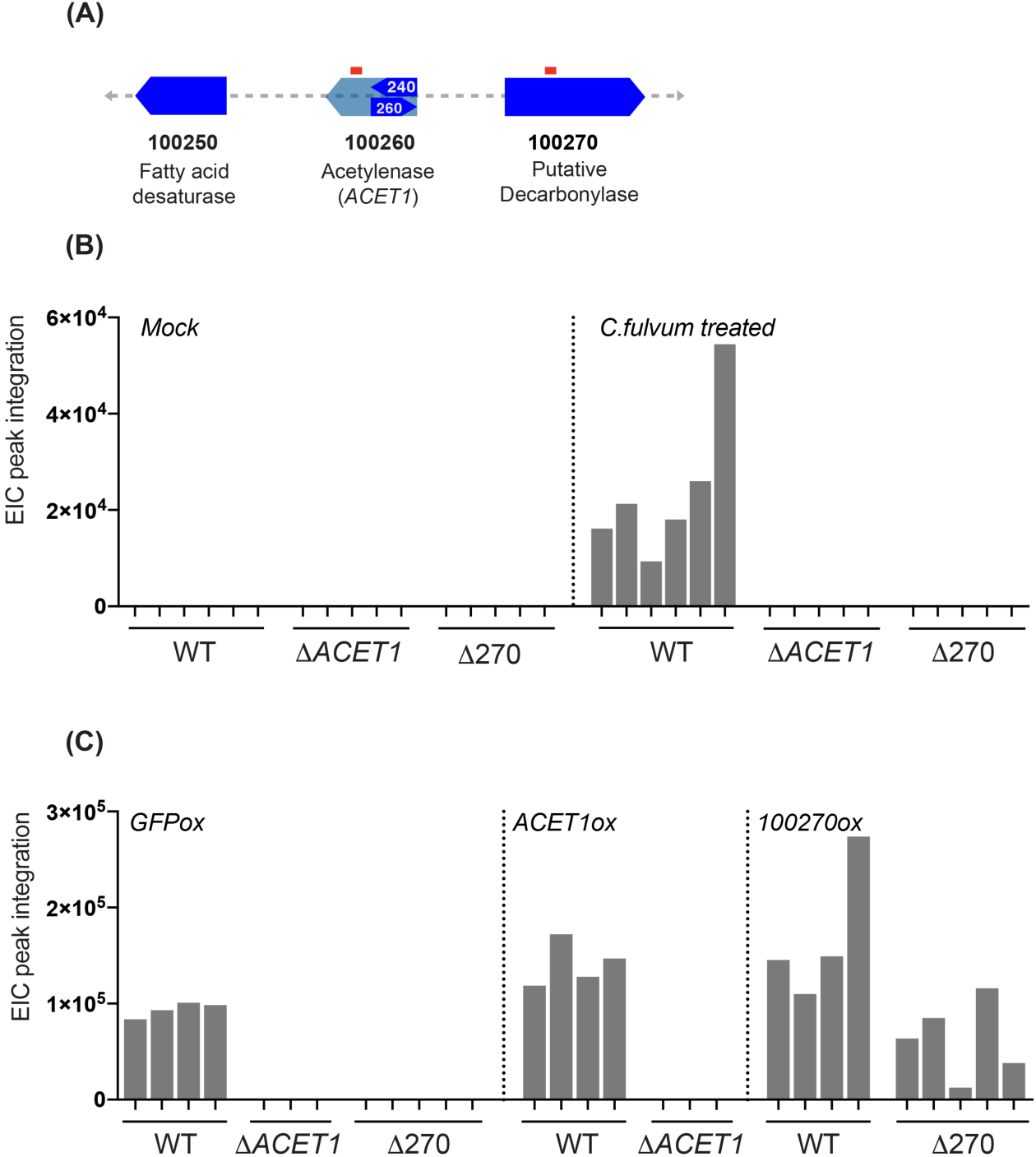
CRISPR/Cas9-induced targeted mutagenesis and gene complementation in tomato. (A) Two CRISPR/Cas9 targeted regions in *ACET1* and 100270 are highlighted in red. (B) Falcarindiol production in wild type (WT, *n*=6) and independent T1 generation CRISPR knock-out mutants of tomato (Δ*ACET1*, *n*=5; Δ270, *n*=5) after Mock or *Cladosporium fulvum* treatment. (C) Falcarindiol production in WT (*n*=4) and mutant lines (Δ*ACET1*, *n*=3; Δ270, *n*=5) after transient overexpression (ox) of genes via *Agrobacterium*-mediated transient assay (*GFPox, ACET1ox,* or *10027ox)*. Genotype of each mutant plant can be found in SI Fig. 12 and the plant ID is # 33, 39, 42, 50, 53 (Δ*ACET1*) and #1, 3, 4, 13, and 20 (Δ270), respectively). Y-axis represents falcarindiol content as determined by integration of EICs from LC- MS data.

Next we performed transient genetic complementation tests for the tomato *ΔACET1* and Δ270 mutants to provide additional evidence that these genes are involved in falcarindiol biosynthesis. We had observed that elicitation of tomato leaves with *A. tumefaciens* is sufficient to induce the falcarindiol biosynthetic pathway (Fig. 3C); therefore, we reasoned that *Agrobacterium-*mediated transient overexpression of wild-type cDNAs for *ACET1* and Solyc12g100270 could both induce the pathway and also provide an enzyme to complement the protein defect present in biosynthetic mutants. We observed that the transient expression of Solyc12g100270 cDNA in Δ270 mutant leaves restores falcarindiol production (Fig. 4C), supporting a direct role for this putative decarbonylase in the biosynthetic pathway. Overexpression of the *ACET1* cDNA did not complement the *ΔACET1* mutants, perhaps due to lack of substrate availability.

Many of the >1400 different acetylenic lipids found in plants are confined to approximately 24 families of higher plants from the Solanales and Apiales orders (*7*). Using the three genes we have connected with falcarindiol biosynthesis in tomato, we investigated whether this gene cluster is conserved among other falcarindiol producers (Fig. S15). Sequence homology indicates high sequence similarity for similar collections of colocalized genes (>80-90% sequence ID at the protein level) in several species of the Solanaceae, including *S. pennellii* (wild tomato), *S. tuberosum* (cultivated potato), and *Capsicum annuum* (bell pepper). Notably, none of these plants have been reported to accumulate falcarindiol. Few genomes from the best known acetylenic fatty acid producers of the Apiales order, including parsley, ginseng, and celery have been sequenced with the exception of carrot. (*30*) In *Daucus carrota*, there are distant orthologs of ACET1 and Solyc12g100250; however, the relatively low sequence identity (<60-70%) at the protein level and lack of gene clustering makes it difficult to predict whether these genes are likely involved in falcarindiol biosynthesis, or if the pathway has arisen in carrot through convergent evolution.

The role of desaturation in the decoration of fatty acids has been associated with changes in membrane fluidity due to alternative structural conformation as well as melting temperature differences imparted by this functionality (*31*), and the presence of multiple acetylenes adds yet another layer of lipid structural tuning that could modify the chemical properties these metabolites. While the biological role of the various chemical modifications on these unusual lipids remains elusive, our work provides new biochemical insights into the metabolic pathways that modify lipids and greatly expands our knowledge of the unusual enzymology that governs their biosynthesis. ACET1, Solyc12g100250 and Solyc12g100270 are predicted to be part of a larger family of non-heme diiron enzymes that include typical desaturases that catalyze selective removal an equivalent of hydrogen from a fatty acid backbone. Similar to the previously characterized acetylenase Crep1, these enzymes appear to have evolved non-canonical function for fatty acid modification.

Our work provides direct biochemical and genetic evidence for the involvement of three core enzymes encoded in a novel cluster of metabolic genes that coordinately act in the biosynthesis of falcarindiol and opens the door for mapping the rich chemistry and biology of highly functionalized oxylipins in plants produced in response to microbial elicitation.

## Acknowledgements

This work was supported by an NIH Genomes to Natural Products U01 GM110699 (to E.S.S.), a USDA NIFA Postdoctoral Fellowship 2016-67012-25102 (to J.E.J.), National Science Foundation IOS-1555957 (to M.B.M.) and generous funding from L’Oreal.

